# Structural basis of DNA targeting by a transposon-encoded CRISPR-Cas system

**DOI:** 10.1101/706143

**Authors:** Tyler S. Halpin-Healy, Sanne E. Klompe, Samuel H. Sternberg, Israel S. Fernández

**Affiliations:** Department of Biochemistry and Molecular Biophysics, Columbia University, New York, NY, 10032, USA

**Author notes:** To whom correspondence should be addressed at (SHS) and (ISF).

## Abstract

Bacteria have evolved adaptive immune systems encoded by Clustered Regularly Interspaced Short Palindromic Repeats (CRISPR) and the CRISPR-associated (Cas) genes to maintain genomic integrity in the face of relentless assault from pathogens and mobile genetic elements [1–3]. Type I CRISPR-Cas systems canonically target foreign DNA for degradation via the joint action of the ribonucleoprotein complex Cascade and the helicase-nuclease Cas3 [4,5] but nuclease-deficient Type I systems lacking Cas3 have been repurposed for RNA-guided transposition by bacterial Tn7-like transposons [6,7]. How CRISPR- and transposon-associated machineries collaborate during DNA targeting and insertion has remained elusive. Here we determined structures of a novel TniQ-Cascade complex encoded by the *Vibrio cholerae* Tn*6677* transposon using single particle electron cryo-microscopy (cryo-EM), revealing the mechanistic basis of this functional coupling. The quality of the cryo-EM maps allowed for de novo modeling and refinement of the transposition protein TniQ, which binds to the Cascade complex as a dimer in a head-to-tail configuration, at the interface formed by Cas6 and Cas7 near the 3’ end of the crRNA. The natural Cas8-Cas5 fusion protein binds the 5’ crRNA handle and contacts the TniQ dimer via a flexible insertion domain. A target DNA-bound structure reveals critical interactions necessary for protospacer adjacent motif (PAM) recognition and R-loop formation. The present work lays the foundation for a structural understanding of how DNA targeting by TniQ-Cascade leads to downstream recruitment of additional transposon-associated proteins, and will guide protein engineering efforts to leverage this system for programmable DNA insertions in genome engineering applications.

---

We previously demonstrated that a transposon derived from *Vibrio cholerae* Tn*6677* undergoes programmable transposition in *E. coli* directed by a CRISPR RNA (crRNA), and that this activity requires four transposon- and three CRISPR-associated genes in addition to a CRISPR array (**Fig. 1a** [7]). Whereas TnsA, TnsB, and TnsC exhibit functions that are consistent with their homologs from a related and well-studied cut-and-paste DNA transposon, *E. coli* Tn7 (reviewed in citePeters:2014aa), we showed that TniQ, a homolog of *E. coli* TnsD, forms a co-complex with the Cascade ribonucleoprotein complex encoded by the Type I-F variant CRISPR-Cas system. This finding suggested an alternative role for TniQ, as compared to the role of *Eco*TnsD in identifying target sites during Tn7 transposition. Rather, we proposed that RNA-guided DNA targeting by Cascade could deliver TniQ to DNA in a manner compatible with downstream transpososome formation, and that TniQ might interact with Cascade near the 3’ end of the crRNA, consistent with RNA-guided DNA insertion occurring approx. 49-bp downstream from the PAM-distal edge of the target site. To determine this unambiguously, we purified the *V. cholerae* TniQ-Cascade complex loaded with a native crRNA and determined its structure by cryo-EM. The overall complex adopts a helical architecture with protuberances at both ends (**Fig. 1** and **Extended Data Fig. 1** and **2**). The global architecture is similar to previously determined structures of Cascade from I-E and I-F systems (**Extended Data Fig. 3**) [8–11] with the exception of a large mass of additional density attributable to TniQ (see below). Maximum likelihood classification methods implemented in Relion3 [12] allowed us to identify significant dynamics in the entire complex, which appears to “breathe”, widening and narrowing the distance between the two protuberances (**Extended Data Fig. 1d** and **Supplementary Movie 1**). The large subunit encoded by a natural Cas8-Cas5 fusion protein (hereafter referred to simply as Cas8) forms one protuberance and recognizes the 5’ end of the crRNA via base- and backbone-specific contacts (**Extended Data Fig. 4, 5a-c, 6a**), akin to the canonical roles played by Cas8 and Cas5 (**Extended Data Fig. 3**). Cas8 exhibits two primary subdomains formed mainly by α-helices, along with a third domain of approximately 100 residues (residues 277 to 385) that is predicted to form three α-helices but could not be built in our maps due to its intrinsic flexibility (**Fig. 1c**). However, low-pass filtered maps revealed that this flexible domain connects with the TniQ protuberance at the opposite end of the crescent-shaped complex (**Extended Data Fig. 2e**). Additionally, there seemed to be a loose coupling between the Cas8 flexible domain and overall “breathing” of the complex, as stronger density for that domain could be observed in the closed state (**Extended Data Fig. 1d** and **Supplementary Movie 1**). Six Cas7 subunits protect much of the crRNA by forming a helical filament along its length (**Fig. 1b** and **d**), similar to other Type I Cascade complexes (**Extended Data Fig. 3** [8–11]. A “finger” motif in Cas7 clamps the crRNA in regular intervals, causing every sixth nucleotide (nt) of the 32-nt spacer to flip out while leaving the flanking nucleotides available for DNA recognition (**Extended Data Fig. 4f**). These bases are pre-ordered in short helical segments, with a conserved phenylalanine stacking below the first base of every segment. Cas7.1, the monomer furthest away from Cas8, interacts with Cas6 (also known as Csy4), which is the ribonuclease responsible for processing of the precursor RNA transcript derived from the CRISPR locus. The Cas6-Cas7.1 interaction is mediated by a β-sheet formed by the contribution of a β-strands from Cas6 and the two β-strands that form the “finger” of Cas7.1 (**Extended Data Fig. 5f**). Cas6 also forms extensive interactions with the conserved stem-loop in the repeat-derived 3’ crRNA handle (**Fig. 1** and **Extended Data Fig. 5d** and **e**), with an arginine-rich α-helix (residues 110 to 128) docked in the major groove, positioning multiple basic residues within interaction distance of the negatively charged RNA backbone. The interaction established between Cas6 and Cas7.1 forms a continuous surface where TniQ is docked, forming the other protuberance of the crescent. The intrinsic flexibility of the complex rendered lower local resolutions in this area of the maps, which we overcame using local alignments masking the area comprising TniQ, Cas6, Cas7.1 and the crRNA handle (**Extended Data Fig. 7**). The enhanced maps allowed for de novo modeling and refinement of TniQ, for which no previous structure or homology model has been reported (**Fig. 2**). Notably, TniQ binds to Cascade as a dimer with head-to-tail configuration (**Fig. 2**), a surprising result given the expectation that *Eco*TnsD functions as a monomer during Tn7 transposition [13]. TniQ is composed of two domains: an N-terminal domain of approximately 100 residues formed by three short α-helices and a second, larger domain of approximately 300 residues with signature sequence for the TniQ family. A DALI search [14] using the refined TniQ model as a probe yielded significant structural similarity of the N-terminal domain to proteins containing Helix-Turn-Helix (HTH) domains (**Extended Data Fig. 8**). This domain is often involved in nucleic acid recognition, however there are reported examples where it has been re-purposed for protein-protein interactions [15]. The remaining C-terminal TniQ-domain is formed by 10 α-helices of variable length and is predicted to contain two tandem zinc finger motifs, though this region was poorly defined in the maps (**Fig. 2**). Overall, the double domain composition of TniQ results in an elongated structure, bent at the junction of the HTH and the TniQ-domain (**Fig. 2**). The HTH domain of one monomer engages the TniQ-domain of the other monomer via interactions between α-helix 3 (H3) and α-helix 11 (H11), respectively, in a tight protein-protein interaction (**Fig. 2c**). This reciprocal interaction is complemented by multiple interactions established between the TniQ-domains from both monomers (up to 45 non-covalent interactions as reported by PISA [16]). Tethering of the TniQ dimer to Cascade is accomplished by specific interactions established with both Cas6 and Cas7. 1 (**Fig. 3**). One monomer of TniQ interacts with Cas6 via its C-terminal TniQ-domain, while the other TniQ monomer contacts Cas7.1 through its N-terminal HTH domain (**Fig. 2b, 3**). The loop connecting alpha-helices H6 and H7 of the TniQ-domain of the first TniQ monomer is inserted in a hydrophobic cavity formed at the interface of two α-helices of Cas6 (**Fig. 3b, d**). The TniQ histidine residue 265 is involved in rearranging the hydrophobic loop connecting H6 and H7 (**Fig. 3d**), which is inserted in the hydrophobic pocket of Cas6 formed by residues L20, Y74, M78, Y83 and F84. The HTH domain of the other TniQ monomer interacts with Cas7.1 through a network of interactions established mainly by α-helix H2 and the linker connecting H2 and H3 (**Fig. 3c, e**). Thus, both the HTH domain and the TniQ-domain exert dual roles to drive TniQ dimerization and dock onto Cascade. In order to explore the structural determinants of DNA recognition by the TniQ-Cascade complex, we determined the structure of the complex bound to a double-stranded DNA (dsDNA) substrate containing the 32-bp target sequence, 5’-CC-3’ PAM, and 20-bp of flanking dsDNA on both ends (**Fig. 4** and **Extended Data Fig. 9**). Density for 28 nucleotides of the target strand (TS) and 8 nucleotides for the non-target strand (NTS) could be confidently assigned in the reconstructed maps (**Fig.4c**). As with previous I-F Cascade structures, Cas8 recognizes the double-stranded PAM within the minor groove (**Extended Data Fig. 10** [10]), and an arginine residue (R246) establishes a stacking interaction with a guanine nucleotide on the TS, which acts like a wedge to separate the double-stranded PAM from the neighboring unwound DNA where base-pairing with the crRNA begins (**Fig. 4b**). Twenty-two nucleotides of the TS within the 32-bp target showed clear density, but surprisingly, the terminal nine nucleotides were not ordered. The TS base-pairs with the spacer region of the crRNA in short, discontinuous, helical segments, as observed previously for I-E and I-F DNA-bound Cascade complexes [10,11] with every 6th base flipped out of the heteroduplex by the insertion of a Cas7 finger (**Extended Data Fig. 6b**). The observed 22-bp heteroduplex is stabilized by the four Cas7 monomers proximal to the PAM (Cas7.6-7.3), but even after local masked refinements, no density could be observed for any TS nucleotides that would base-pair with the 3’ end of the crRNA spacer bound by Cas7.2 and Cas7.1. These two Cas7 monomers are proximal to Cas6 and in the region previously described to exhibit dynamics due to the interaction of the Cas8 flexible domain with the inner face of the TniQ-dimer. In addition, the disordered nucleotides also correspond to positions 25-28 of the target site where RNA-DNA mismatches are detrimental for RNA-guided DNA integration [7]. Thus, we propose the possibility that the partial R-loop structure we observed may represent an intermediate conformation refractory to integration, and that further structural rearrangements may be critical for further stabilization of an open conformation, possibly driven by recruitment of the TnsC ATPase. Here we present the first cryo-EM structures of a CRISPR-Cas effector complex bound to the transposition protein TniQ, with and without target DNA. These structures reveal the unexpected presence of TniQ as a dimer that forms bipartite interactions with Cas6 and Cas7.1 within the Cascade complex, forming a likely recruitment platform for downstream-acting transposition proteins18 (**Fig. 4d**). Our structures furthermore reveal a possible fidelity checkpoint, whereby formation of a complete R-loop requires conformational rearrangements that may depend on extensive RNA-DNA complementarity and/or downstream factor recruitment; this proofreading step could account for the highly specific RNA-guided DNA integration we previously reported for the *V. cholerae* transposon [7]. In light of recent work demonstrating exaptation of Type V-K CRISPR-Cas systems by similar Tn7-like transposons that also encode TniQ [17,18], it will be interesting to determine whether tethering of TniQ to evolutionarily distinct CRISPR RNA effector complexes - Cascade or Cas12k - is a general theme of RNA-guided transposition.

**Fig.1.**
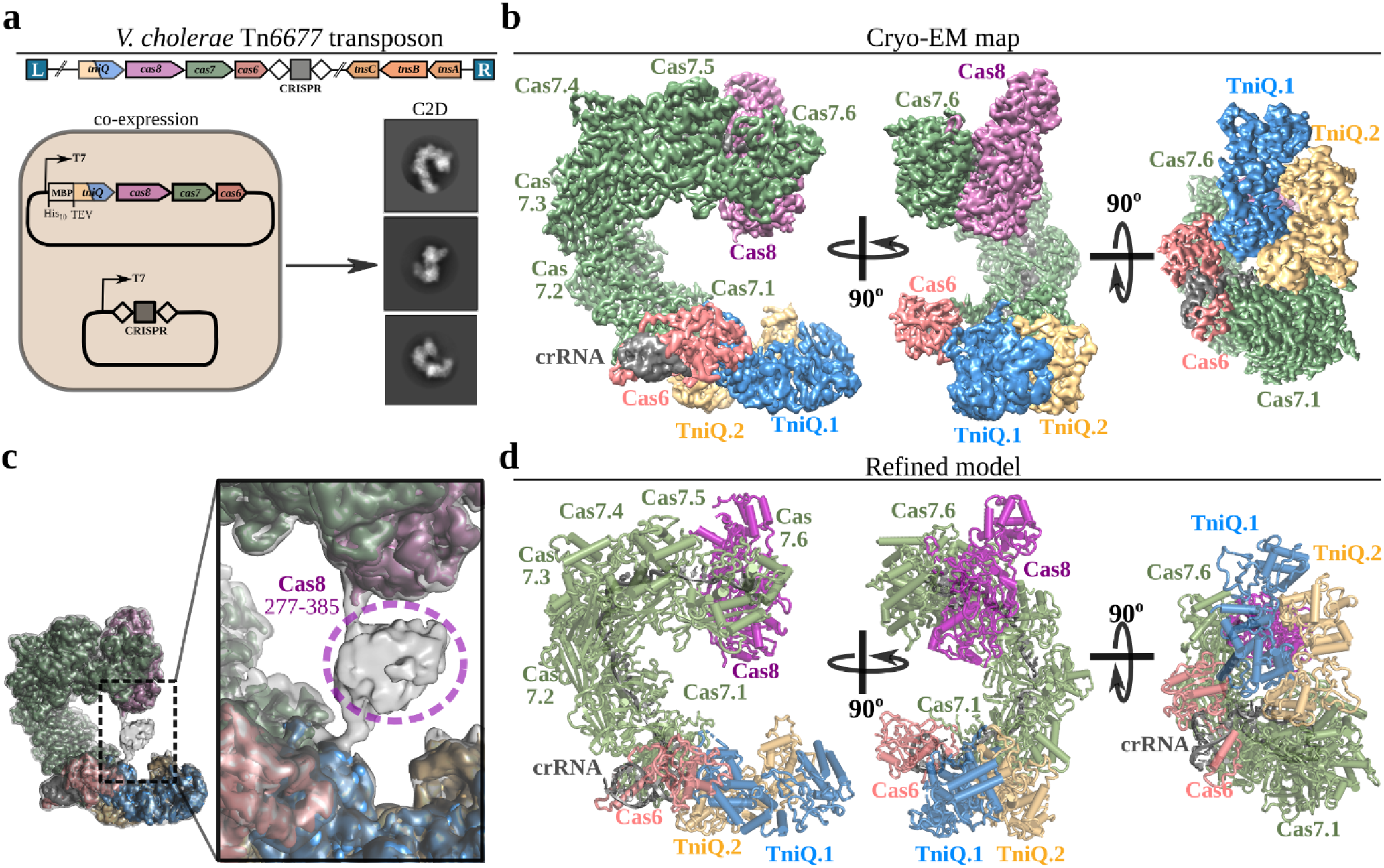
Overall architecture of the *V. cholerae* TniQ-Cascade complex. **a**, Genetic architecture of the Tn*6677* transposon (top), and plasmid constructs used to express and purify the TniQ-Cascade co-complex. Selected cryo-EM reference-free 2D classes in multiple orientations are shown on the right. **b**, Orthogonal views of the cryo-EM map for the TniQ-Cascade complex, showing Cas8 (pink), six Cas7 monomers (green), Cas6 (salmon), crRNA (grey), and TniQ monomers (blue, yellow). The complex adopts a helical architecture with protuberances at both ends. **c**, A flexible domain in Cas8 comprising residues 277-385 (grey) could only be visualized in low-pass filtered maps. The unsharpened map is shown as semi-transparent, grey map overlaid on the post-processed map segmented and colored according to a. **d**, Refined model for the TniQ-Cascade complex derived from the cryo-EM maps shown in **b**.

**Fig.2.**
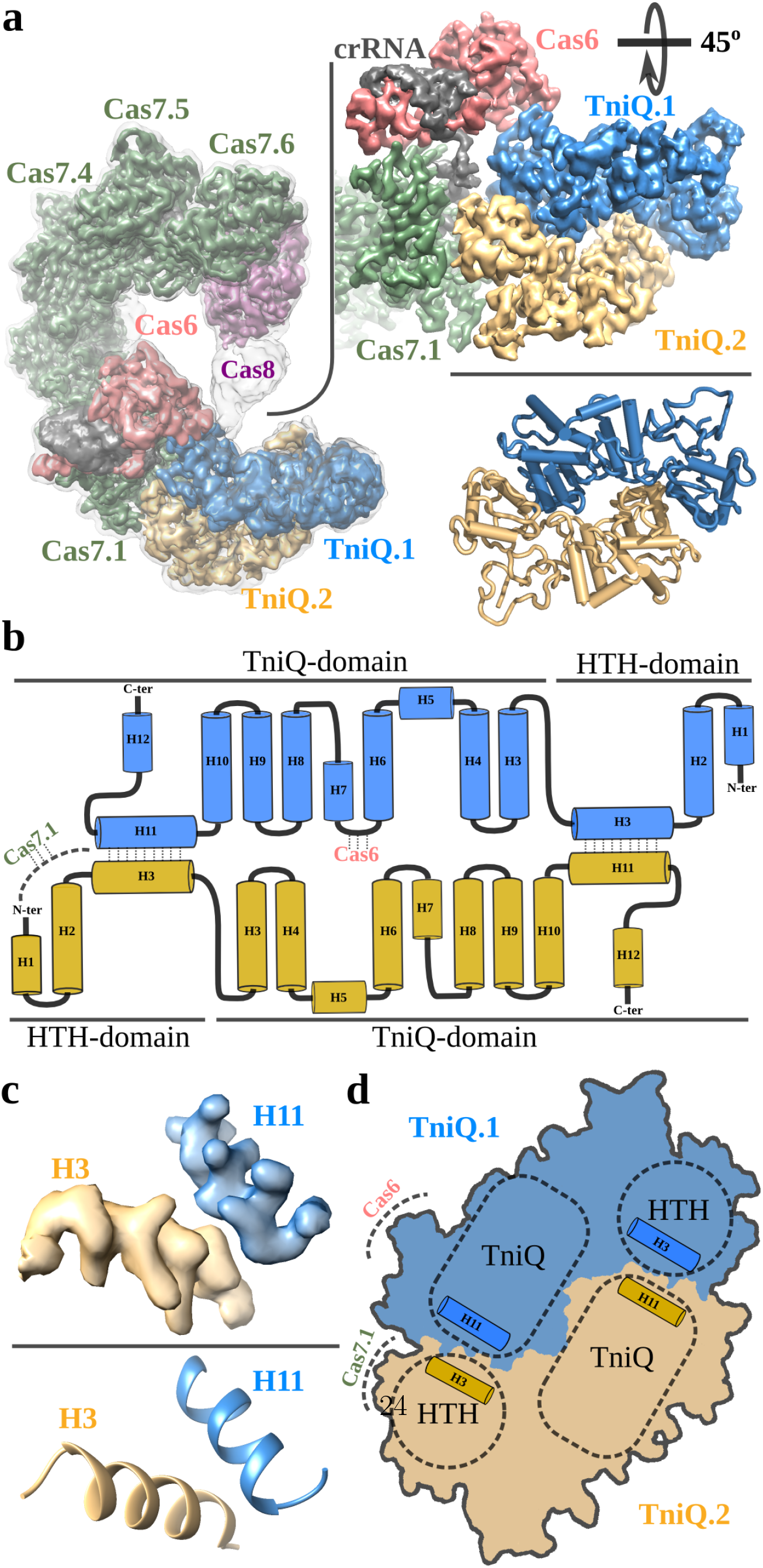
TniQ binds Cascade in a dimeric, head-to-tail configuration. **a**, Left, overall view of the TniQ-Cascade cryo-EM unsharpened map (grey) overlaid on the post-processed map segmented and colored as in **Fig.1**. Right, cryo-EM map (top) and refined model (bottom) of the TniQ dimer. The two monomers interact with each other in a head-to-tail configuration and are anchored to Cascade via Cas6 and Cas7.1. **b**, Secondary structure diagram of the TniQ dimer: eleven α-helices are organized into an N-terminal Helix-Turn-Helix (HTH) domain and a C-terminal TniQ-domain. Dimer interactions between H3 and H11 are indicated, as are interaction sites with Cas6 and Cas7.1. **c**, Cryo-EM density for the H3-H11 interaction shows clear side-chain features (top), allowing accurate modeling of the interaction (bottom). **d**, Schematic of the dimer interaction, showing the important dimerization interface between the HTH and TniQ-domain.

**Fig.3.**
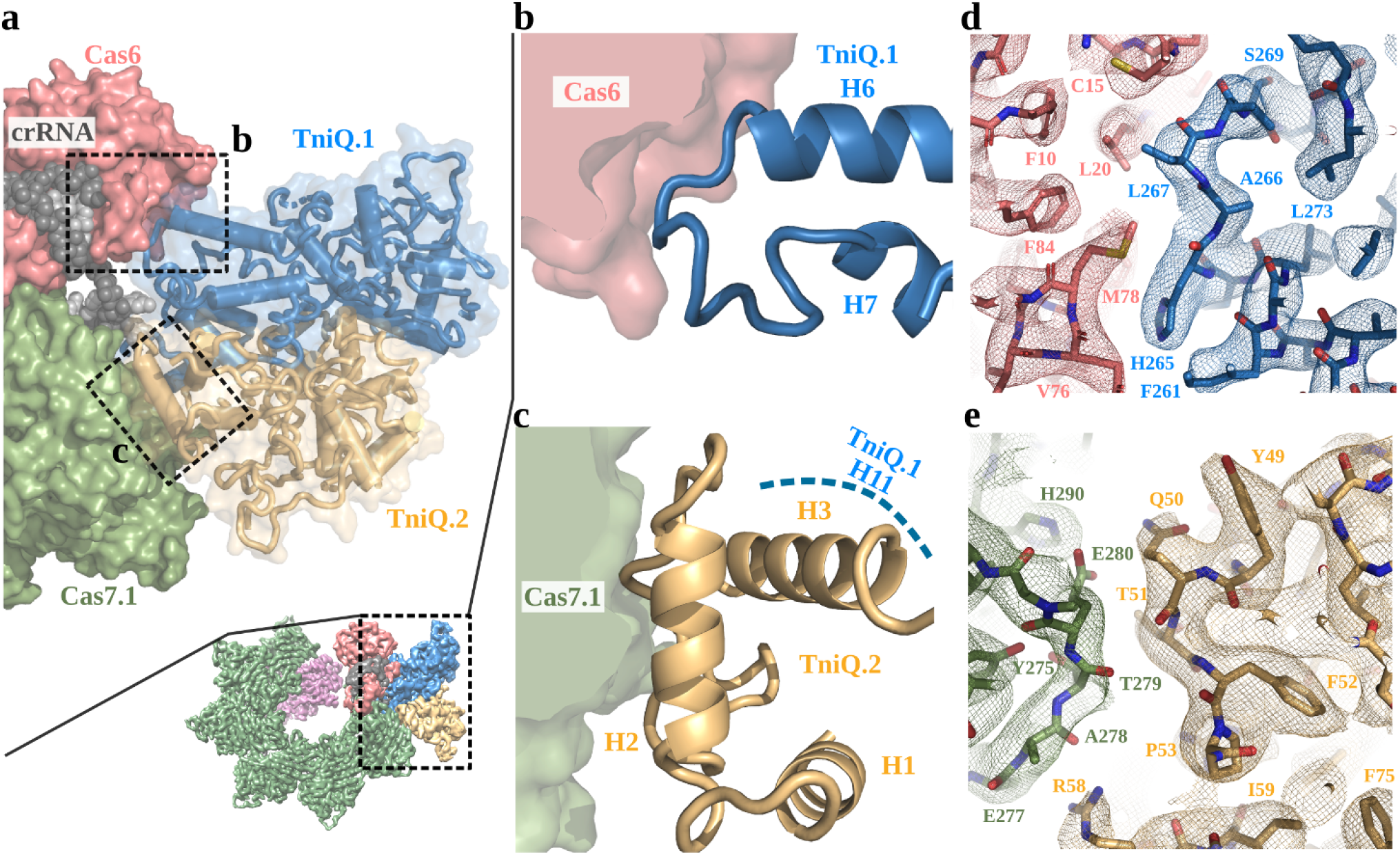
Cas6 and Cas7.1 form a binding platform for TniQ. **a**, Top, zoomed area showing the interaction site of Cascade and the TniQ dimer. Cas6 and Cas7.1 are displayed as molecular Van der Waals surfaces, the crRNA is shown as grey spheres, and the TniQ monomers as ribbons. **b**, The loop connecting TniQ.1 α-helices H6 and H7 (blue) binds within a hydrophobic cavity of Cas6. **c**, Cas7.1 interacts via with the HTH domain of the TniQ.2 monomer (yellow), mainly through H2 and the loop connecting H2 and H3. **d-e**, Experimental cryo-EM densities observed for the TniQ-Cas6 (textbfd) and TniQ-Cas7.1 (textbfe) interaction.

**Fig.4.**
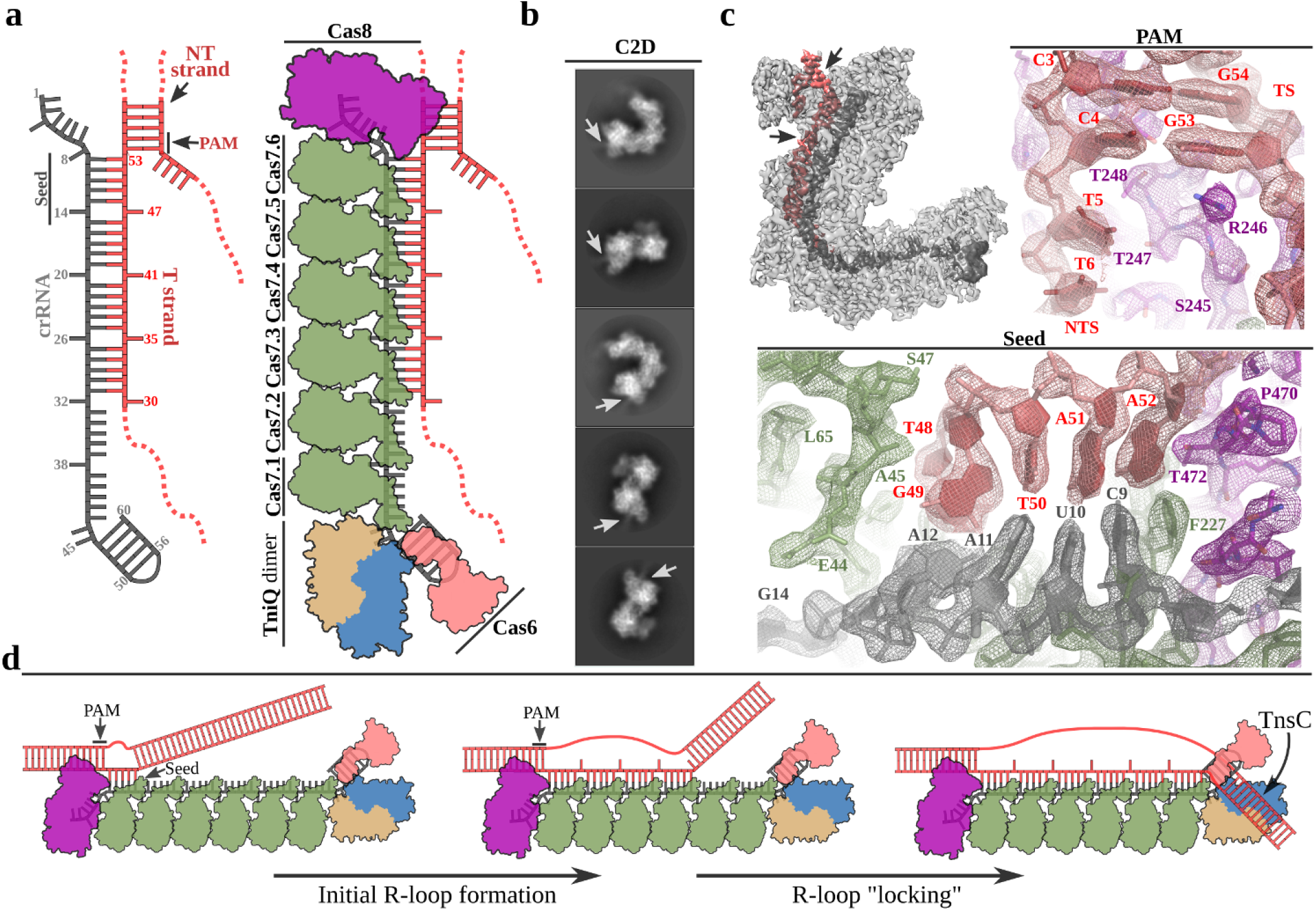
DNA-bound structure of the TniQ-Cascade complex. **a**, Schematic of crRNA and the portion of the dsDNA substrate that was experimentally observed within the electron density map for DNA-bound TniQ-Cascade. Target Strand (TS), non-target strand (NTS), as well as the PAM and seed regions are indicated. **b**, Selected cryo-EM reference-free 2D classes for DNA-bound TniQ-Cascade; density corresponding to dsDNA could be directly observed protruding from the Cas8 component in the 2D averages (white arrows). **c**, Cryo-EM map for DNA-bound TniQ-Cascade. The crRNA is in dark grey and the DNA is in red. On the right and bottom, detailed views for the PAM and seed recognition regions of the map, with refined models represented as sticks within the electron density. Cas8 is shown in pink, Cas7 in green, crRNA in grey, and DNA in red. **d**, The *V. cholerae* transposon encodes a TniQ-Cascade co-complex that utilizes the sequence content of the crRNA to bind complementary DNA target sites (left). We propose that the incomplete R-loop observed in our structure (middle) represents an intermediate state that may precede a downstream “locking” step involving proofreading of the RNA-DNA complementarity. TniQ is positioned at the PAM-distal end of the DNA-bound Cascade complex, where it likely interacts with TnsC during downstream steps of RNA-guided DNA insertion.

## Methods

### TniQ-Cascade purification

Protein components of TniQ-Cascade were expressed from a pET-derivative vector containing the native *V. cholerae* tniQ-cas8-cas7-cas6 operon with an N-terminal His10-MBP-TEVsite fusion on TniQ. The crRNA was expressed separately from a pACYC-derivative vector containing a minimalrepeat-spacer-repeat CRISPR array encoding a spacer from the endogenous *V. cholerae* CRISPR array. The TniQ-Cascade complex was overexpressed and purified as described previously [7], and was stored in Cascade Storage Buffer (20 mM Tris-Cl, pH 7.5, 200 mM NaCl, 1 mM DTT, 5% glycerol).

### Sample preparation for electron microscopy

For negative staining, 3 μL of purified TniQ-Cascade ranging from 100 nM to 2 μM was incubated with plasma treated (H2/O2 gas mix, Gatan Solarus) CF400 carbon-coated grids (EMS) for 1 minute. Excess solution was blotted and 3 μL of 0.75% uranyl formate was added for an additional minute. Excess stain was blotted away and grids were air-dried overnight. Grid screening for both negative staining and cryo conditions was performed on a Tecnai-F20 microscope (FEI) operated at 200 KeV and equipped with a Gatan K2-Summit direct detector. Microscope operation and data collection were carried out using the Leginon/Appion software. Initial negative staining grid screening allowed determination of a suitable concentration range for cryo conditions. Several grid geometries were tested in the 1-4 μM concentration range for cryo conditions using a Vitrobot Mark-II operated at 4 C, 100% humidity, blot force 3, drain time 0, waiting time 15 seconds, and blotting times ranging from 3-5 seconds. The best ice distribution and particle density was obtained with 0.6/1 UltrAuFoil grids (Quantifoil).

### Electron microscopy

A preliminary dataset of 300 images in cryo was collected with the Tecnai-F20 microscope using a pixel size of 1.22 Å/pixel with illumination conditions adjusted to 8 e-/pixel/second with a frame window of 200 ms. Preprocessing and image processing were integrally done in Relion3 [12] with ctf estimation integrated via a wrapper to Gctf [19]. An initial model computed using the SGD algorithm [20] implemented in Relion3 was used as initial reference for a refine 3D job that generated a sub-nanometric reconstruction with approximately 10,000 selected particles. Clear secondary structure features in the 2D averages and the 3D reconstruction could be identified. For the DNA-bound TniQ-Cascade complex containing DNA, we pre-incubated two complementary 74-nt oligonucleotides (NTS: 5’TTCATCAAGCCATTGGACCGCCTTACAGGACGCTTTGGCTTCATTGCTTTTCAGCTTCGCCTTGACGGCCAAAA-3’, TS: 5’TTTTGGCCGTCAAGGCGAAGCTGAAAAGCAATGAAGCCAAAGCGTCCTGTAAGGCGGTCCAATGGCTTGATGAA-3’) for 5 minutes at 95° C in hybridization buffer (20 mM Tris-Cl, pH 7.5, 100 mM KCl, 5 mM MgCl2) to form dsDNA, which was subsequently aliquoted and flash frozen. Complex formation was performed by incubating a 3x molar excess of dsDNA with TniQ-Cascade at 37° C for 5 minutes prior to vitrification, which followed the conditions optimized for the apo complex (defined as TniQ-Cascade with crRNA but no DNA ligand). High resolution data for the apo complex were collected in a Tecnai-Polara-F30 microscope operated at 300 KeV equipped with a K3 direct detector (Gatan). A 30 μm C2 aperture was used with a pixel size of 0.95 Å/pixel and illumination conditions in microprobe mode adjusted to a fluence of 16e-/pixel/second. Four-second images with a frame width of 100 ms (1.77 e-/2/frame) were collected in counting mode. For the DNA-bound complex, high resolution data were collected in a Titan Krios microscope (FEI) equipped with an energy filter (20 eV slit width) and a K2 direct detector (Gatan) operated at 300 KeV. A 50 μm C2 aperture was used with a pixel size of 1.06 Å/pixel and illumination conditions adjusted in nanoprobe mode to a fluence of 8e-/pixel/second. Eight-second images with a frame width of 200 ms (1.42 e-/2/frame) were collected in counting mode.

### Image processing

Motion correction was performed for every micrograph applying the algorithm described for Mo-tioncor2 [21] implemented in Relion3 with 5 by 5 patches for the K2 data and 7 by 5 patches for the K3 data. Parameters of the contrast transfer function for each motion-corrected micrograph were obtained using Gctf integrated in Relion3. Initial particle picking of a subset of 200 images randomly chosen was performed with the Laplacian tool of the Auto-picking module of Relion3, using an estimated size for the complex of 200 Å15,000 particles were extracted in a 300 pixels box size and binned 3 times for an initial 2D classification job. Selected 2D averages from this job were used as templates for Auto-picking of the full dataset. The full dataset of binned particles was subjected to a 2D classification job to identify particles able to generate averages with clear secondary structure features. The selected subgroup of binned particles after the 2D classification selection was refined against a 3D volume obtained by SGD with the F20 data. This “consensus” volume was inspected to localize areas of heterogeneity which were clearly identified at both ends of the crescent shape characteristic of this complex. Both ends were then individually masked using soft masks of around 20 pixels that were subsequently used in classification jobs without alignments in Relion3. The T parameter used for this classification job was 6 and the total number of classes was 10. This strategy allowed us to identify two main population of particles which correspond to an “open” and “closed” state of the complex. Particles from both subgroups were separately re-extracted to obtain unbinned datasets for further refinement. New features implemented in Relion3, namely Bayesian polishing and ctf parameters refinement, allowed the extension of the resolution to 3.4, 3.5 and 2.9 Å for the two apo and the DNA-bound complexes, respectively. Post processing was performed with a soft-mask of 5 pixels being the B-factor estimated automatically in Relion3 following standard practice. A final set of local refinements was performed with the masks used for classification. The locally aligned maps exhibit very good quality for the ends of the C-shape. These maps were used for de novo modeling and initial model refinement.

### Model building and refinement

For the Cas7 and Cas6 monomers, the *E. coli* homologs (PDB accession code 4TVX) were initially docked with Chimera [22] and transformed to poly-alanine models. Substantial rearrangement of the finger region of Cas7 monomers, as well as other secondary structure elements of Cas6, were performed manually in COOT [23] before amino acid substitution of the poly-alanine model. Well-defined bulky side chains of aromatic residues allowed a confident assignment of the register. The crRNA was also well defined in the maps and was traced de novo with COOT. For Cas8 and TniQ in particular, no structural similarity was found in the published structures able to explain our densities. Locally refined maps using soft masks at both ends of the crescent-shaped complex rendered well-defined maps below 3.5 Å resolution. These maps were used for manual de novo tracing of a poly-alanine model in COOT that was subsequently mutated to the *V. cholerae* sequences. Bulky side chains for aromatic residues showed excellent density and were used as landmarks to adjust the register of the sequence. For refinement, an initial step of real space refinement against the cryo-EM maps was performed with the phenix real space refinement tool of the Phenix package [24], with secondary structure restraints activated. A second step of reciprocal space refinement was performed in Refmac5 [25], with secondary restraints calculated with Prosmart [26] and LibG [27]. Weight of the geometry term versus the experimental term was adjusted to avoid overfitting of the model into cryo-EM map, as previously reported30. Model validation was performed in Molprobity [28].

## Data availability

Maps and models have been deposited in the EMDB (accession codes 20349, 20350 and 20351) and the PDB (accession codes 6PIF, 6PIG and 6PIJ).

## Acknowledgements

We acknowledge Bob Grassucci and Zhening Zhang for technical assistance in cryo-EM data acquisition. Part of this work was performed at the Simons Electron Microscopy Center and National Resource for Automated Molecular Microscopy located at the New York Structural Biology Center, supported by grants from the Simons Foundation (SF349247), NYSTAR, and the NIH National Institute of General Medical Sciences (GM103310).

## EXTENDED DATA FIGURES

**Extended Data Fig. 1.**
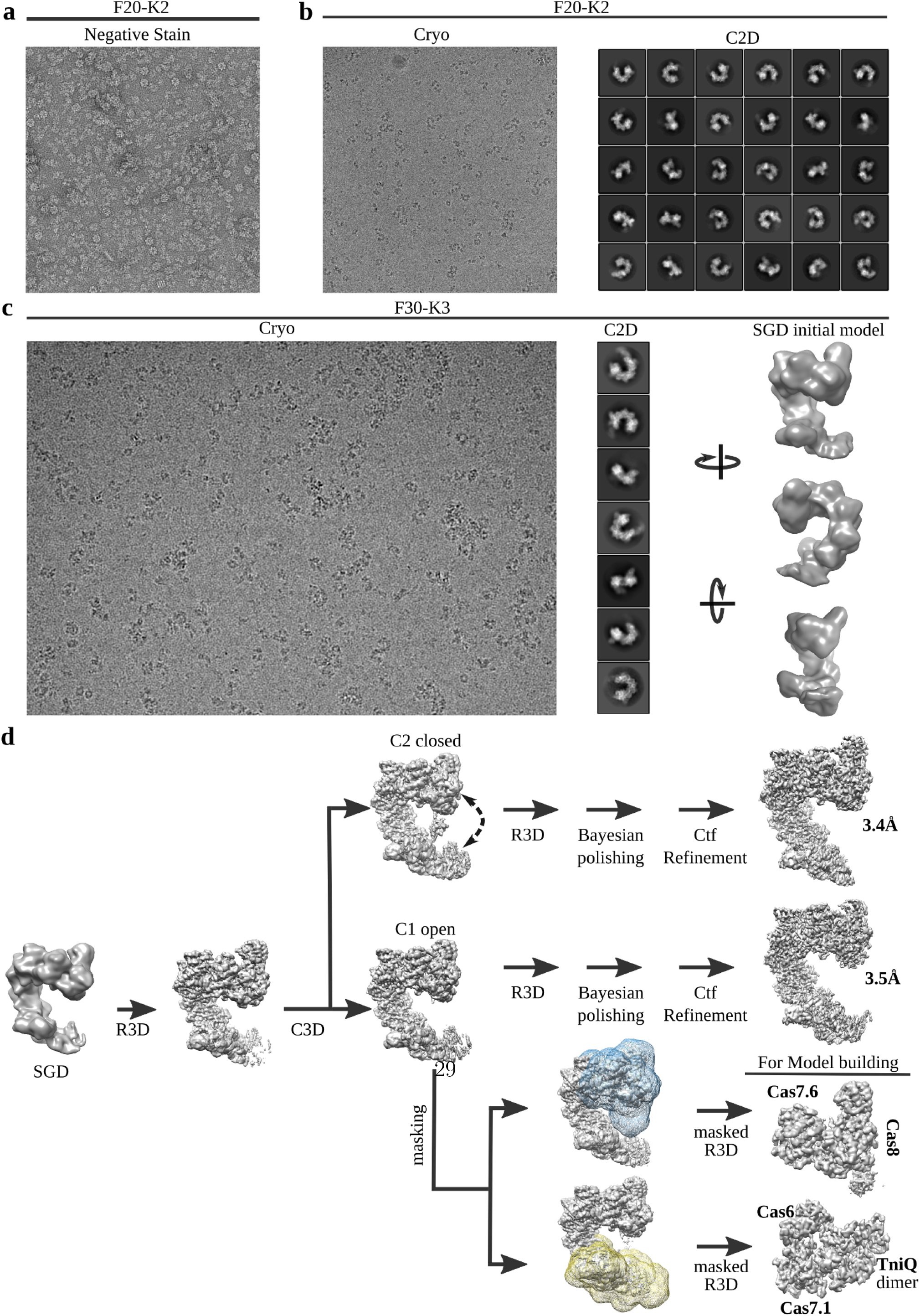
Cryo-EM sample optimization and image processing workflow. **a**, Representative negatively stained micrograph for 500 nM TniQ-Cascade. **b**, Left, representative cryo-EM image for 2 μM TniQ-Cascade. A small dataset of 200 images was collected in a Tecnai-F20 microscope equipped with a Gatan K2 camera. Right, reference-free 2D class averages for this initial cryo-EM dataset. **c**, Left, representative image from a large dataset collected in a Tecnai Polara microscope equipped with a Gatan K3 detector. Middle, detailed 2D class averages were obtained that were used for initial model generation using the SGD algorithm implemented in Relion3 (right). **d**, Image processing workflow used to identify the two main classes of the TniQ-cascade complex in open and closed conformations. Local refinements with soft masks were used to improve the quality of the map within the terminal protuberances of the complex. These maps were instrumental for *de novo* modeling and initial model refinement.

**Extended Data Fig. 2.**
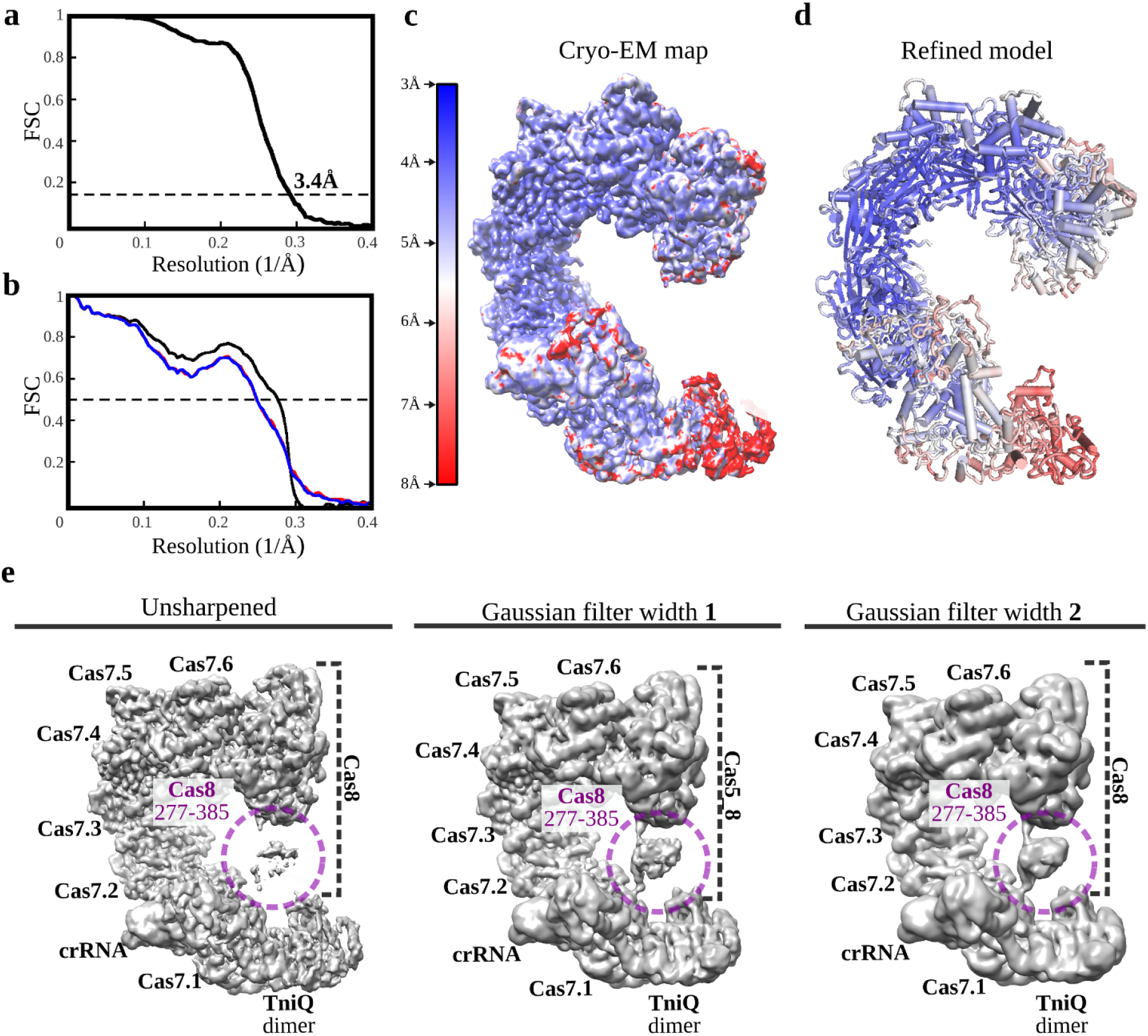
Fourier Shell Correlation (FSC) curves, local resolution, and unsharpened filter maps for the TniQ-Cascade complex in closed conformation. **a**, Gold-standard FSC curve using half maps; the global resolution estimation is 3.4 Åby the FSC 0.143 criterion. **b**, Cross-validation model-vs-map FSC. Blue curve, FSC between the shacked model refined against half map 1; red curve, FSC against half map 2, not included in the refinement; black curve, FSC between final model against the final map. The overlap observed between the blue and red curves guarantees a non-overfitted model [29]. **c**, Unsharpened map colored according to local resolutions, as reported by RESMAP [30]. **d**, Final model colored according to B-factors calculated by REFMAC. **e**, A flexible Cas8 domain encompassing residues 277-385 contacts the TniQ dimer at the other side of the crescent shape. Applying a Gaussian filter of increasing width to the unsharpened map allows for a better visualization of this flexible region.

**Extended Data Fig. 3.**
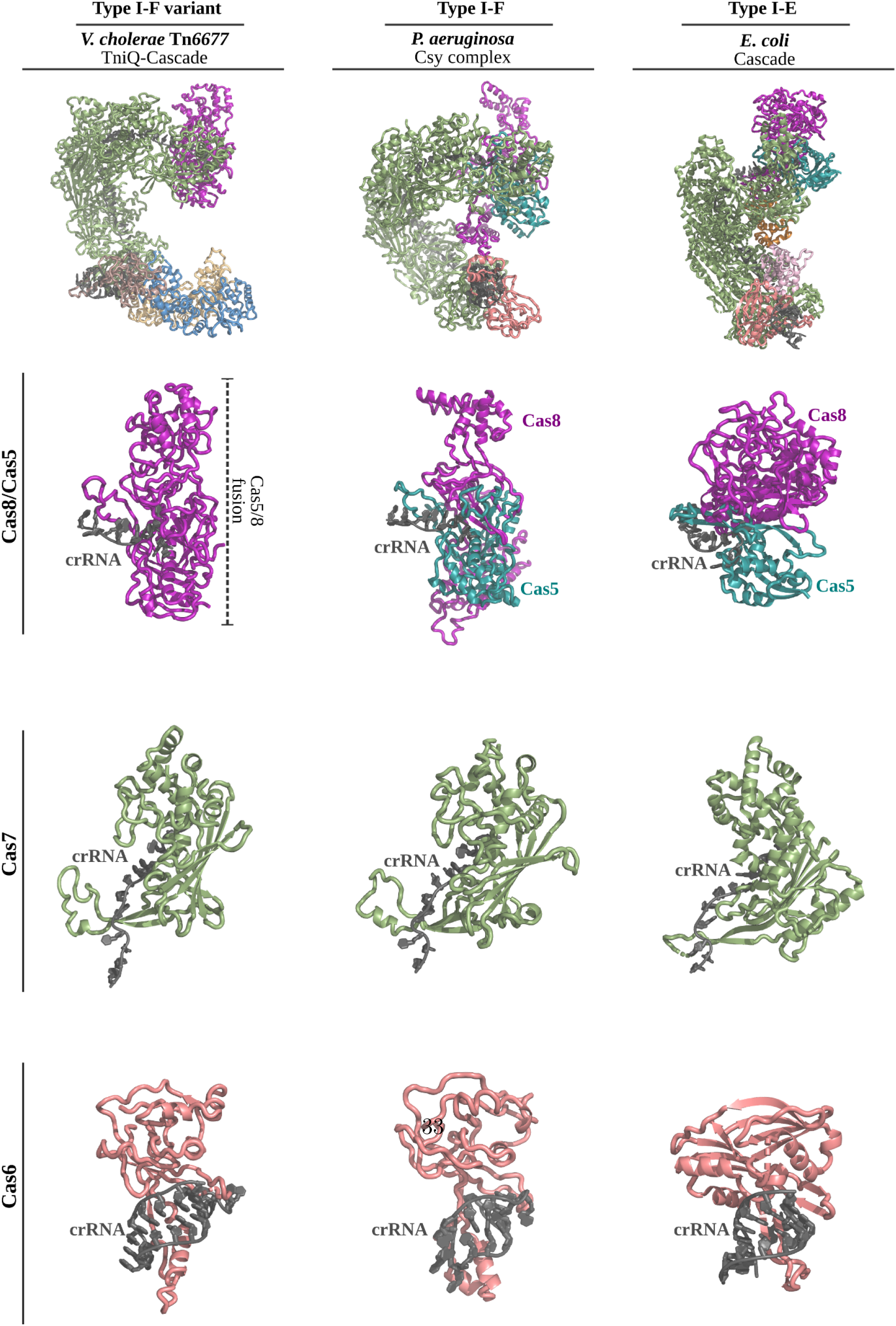
Superposition of TniQ-Cascade with structurally similar Cascade complexes. The *V. cholerae* I-F variant TniQ-Cascade complex (left) was superposed with *Pseudomonas aeruginosa* I-F Cascade11 (also known as Csy complex; middle, PDB ID: 6B45) and *Escherichia coli* I-E Cascade9 (right, PDB ID: 4TVX). Shown are superpositions of the entire complex (top), the Cas8 and Cas5 subunits with the 5’ crRNA handle (middle top), the Cas7 subunit with a fragment of crRNA (middle bottom), and the Cas6 subunit with the 3’ crRNA handle (bottom).

**Extended Data Fig. 4.**
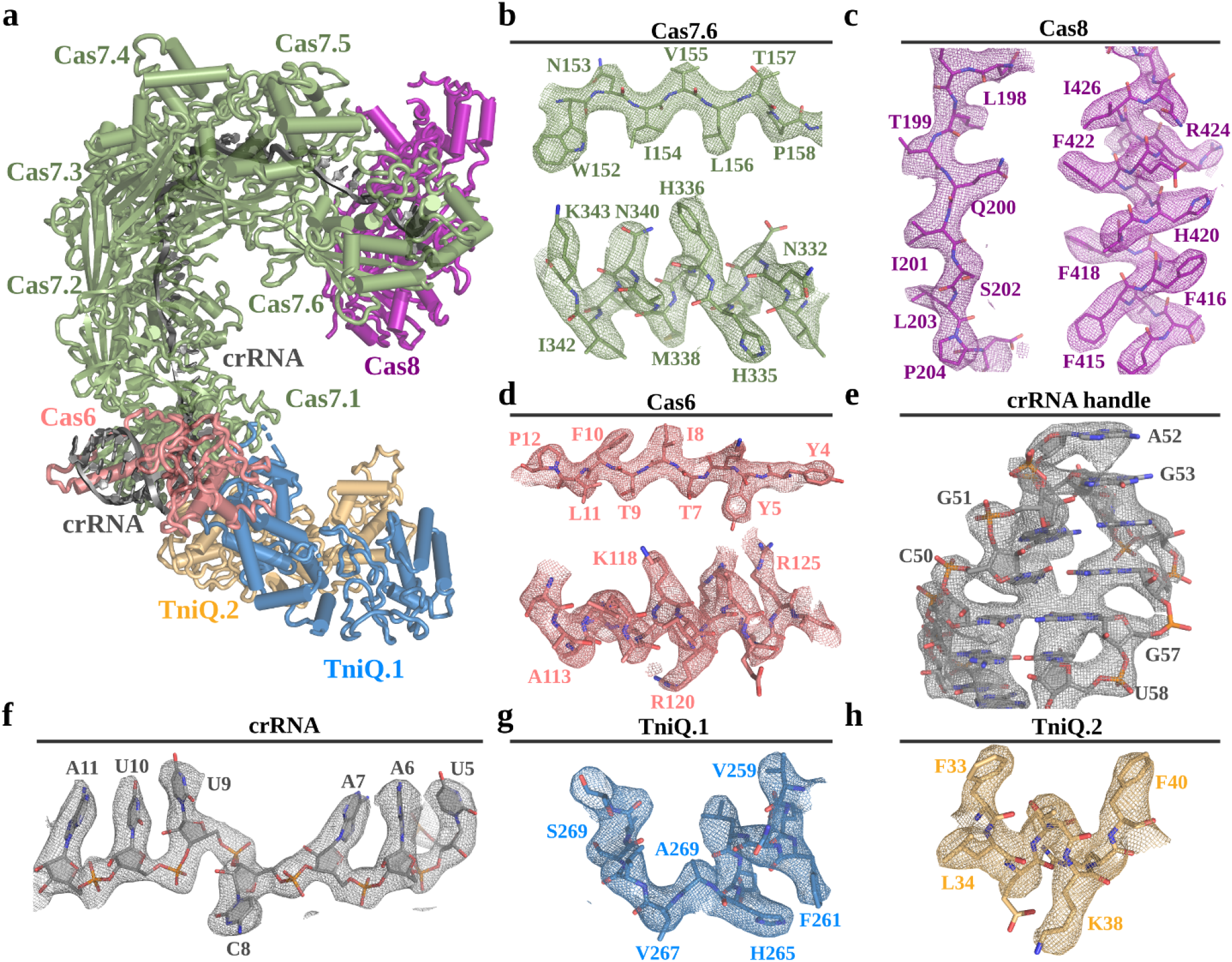
Representative cryo-EM densities for all the components of the TniQ-Cascade complex in closed conformation. **a**, Final refined model of TniQ-Cascade, with Cas8 in purple, Cas7 monomers in green, Cas6 in red, the TniQ monomers in blue and yellow, and the crRNA in grey. **b-h**, Final refined model inserted in the final cryo-EM density for select regions of all the molecular components of the TniQ-Cascade complex. Residues are numbered.

**Extended Data Fig. 5.**
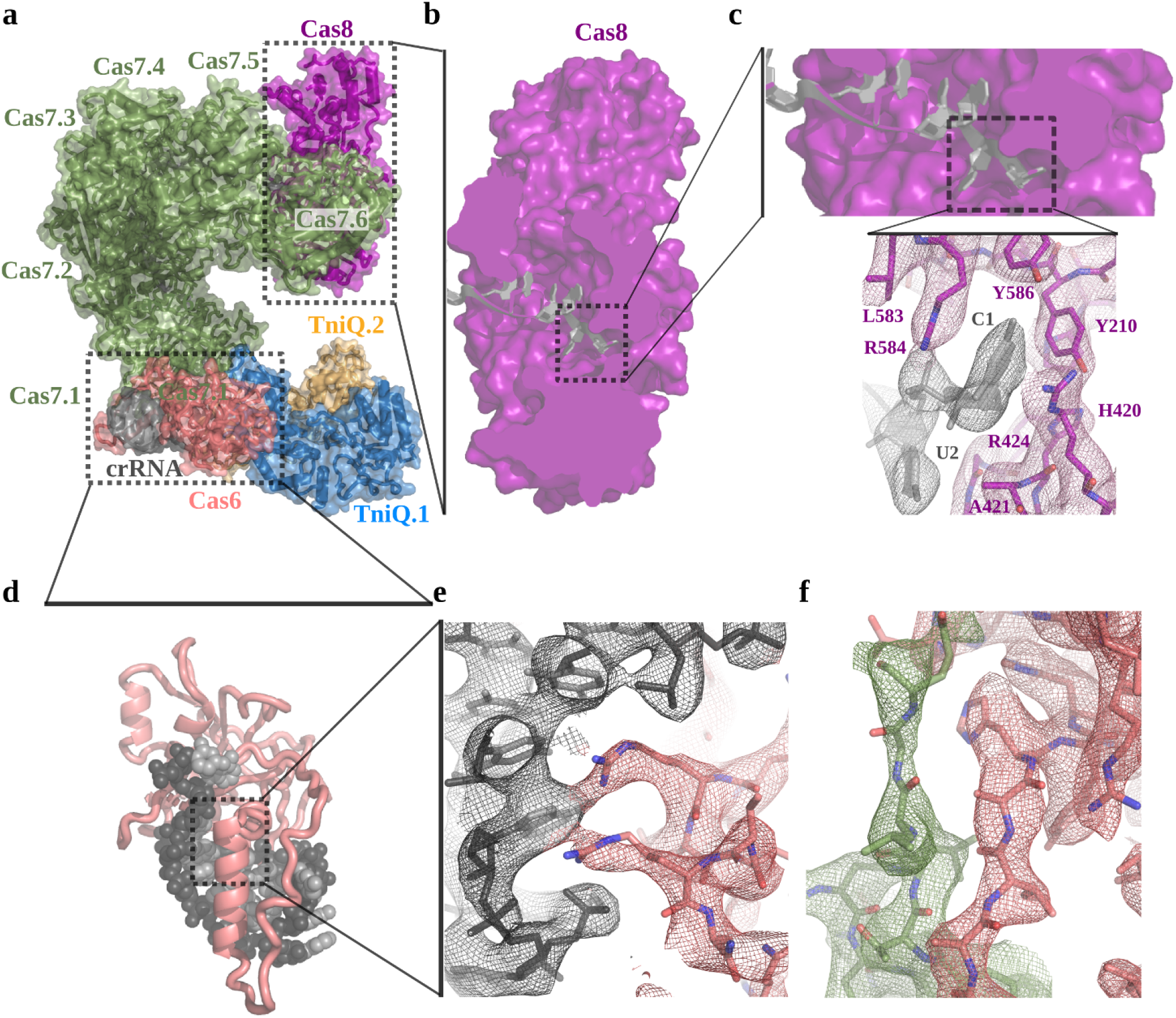
Cas8 and Cas6 interaction with the crRNA. **a**, Refined model for the TniQ-Cascade shown as ribbons inserted in the semitransparent Van der Walls surface, colored as in **Fig 1. b** and **c**, Zoomed view of Cas8, which interacts with the 5’end of the crRNA. The inset shows electron density for the highlighted region, where the base of nucleotide C1 is stabilized by stacking interactions with arginine residues R584 and R424. **d**, Cas6 interacts with the 3’ end of the crRNA “handle” (nucleotides 45-60). **e**, An arginine-rich α-helix is deeply inserted within the major groove of the terminal stem-loop. This interaction is mediated by electrostatic interactions between basic residues of Cas6 and the negatively charged phosphate backbone of the crRNA. **f**, Cas6 (red) also interacts with Cas7.1 (green), establishing a β-sheet formed by β-strands contributed from both proteins.

**Extended Data Fig. 6.**
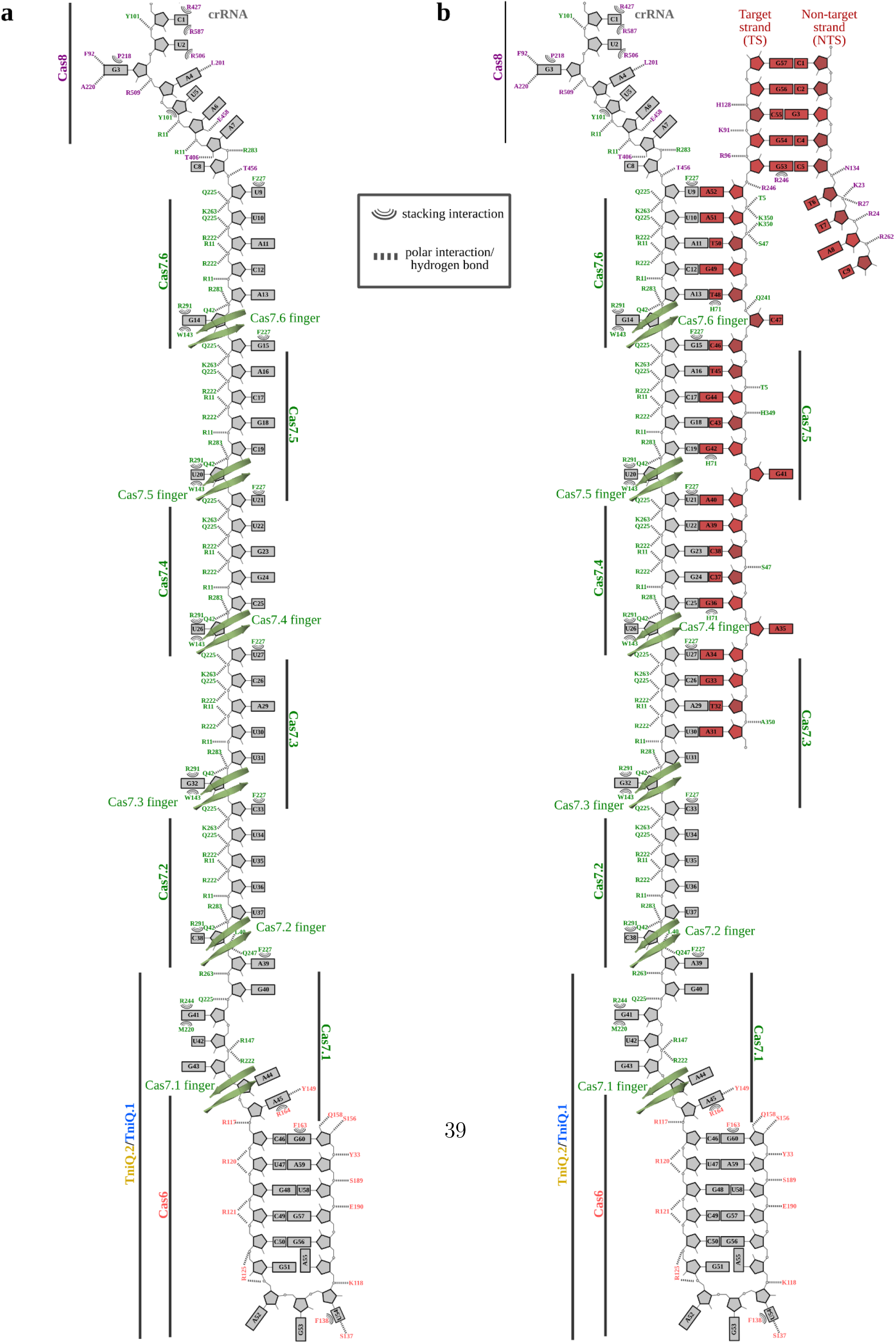
Schematic representation of crRNA and target DNA recognition by TniQ-Cascade. **a**, TniQ-Cascade residues that interact with the crRNA are indicated. Approximate location for all protein components of the complex are also shown, as well as the position of each Cas7 “finger”.? **b**, TniQ-Cascade residues that interact with crRNA and target DNA, shown as in **a**.

**Extended Data Fig. 7.**
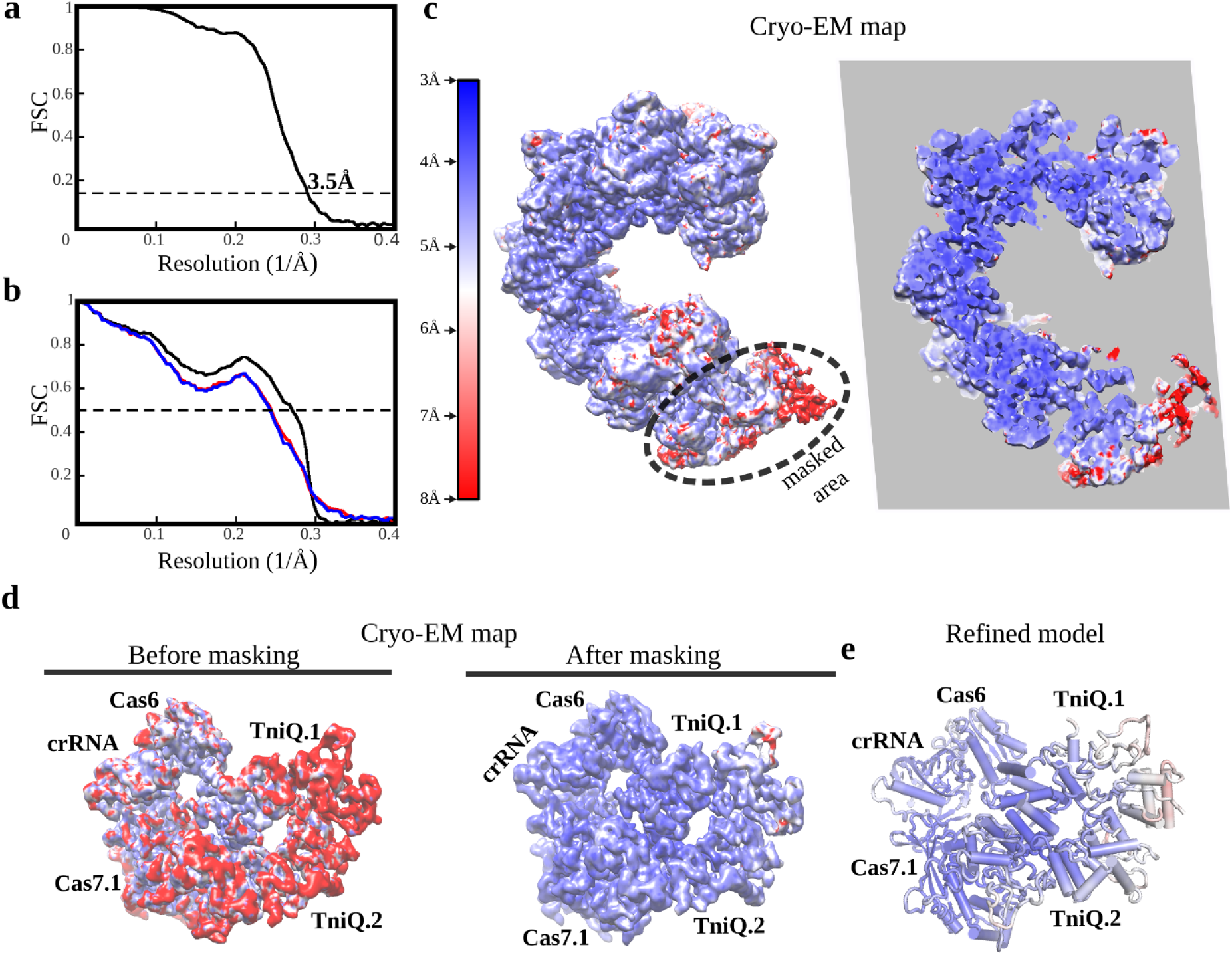
Fourier Shell Correlation (FSC) curves, local resolution, and local refined maps for the TniQ-Cascade complex in open conformation. **a**, Gold-standard FSC curve using half maps; the global resolution estimation is 3.5 Å by the FSC 0.143 criterion. **b**, Cross-validation model-vs-map FSC. Blue curve, FSC between shacked model refined against half map 1; red curve, FSC against half map 2, not included in the refinement; black curve, FSC between final model against the final map. The overlapping between the blue and red curves guarantees a non-overfitted model. **c**, Unsharpened map colored according to local resolutions, as reported by RESMAP. Right, slice through the map shown on the left. **d**, Local refinements with soft masks improved the maps in flexible regions. Shown the region of the map corresponding to the TniQ dimer. Unsharpened maps colored according to the local resolution estimations are shown before (left) and after (right) masked refinements. **e**, Final model for the TniQ dimer region, colored according to the local B-factors calculated by REFMAC.

**Extended Data Fig. 8.**
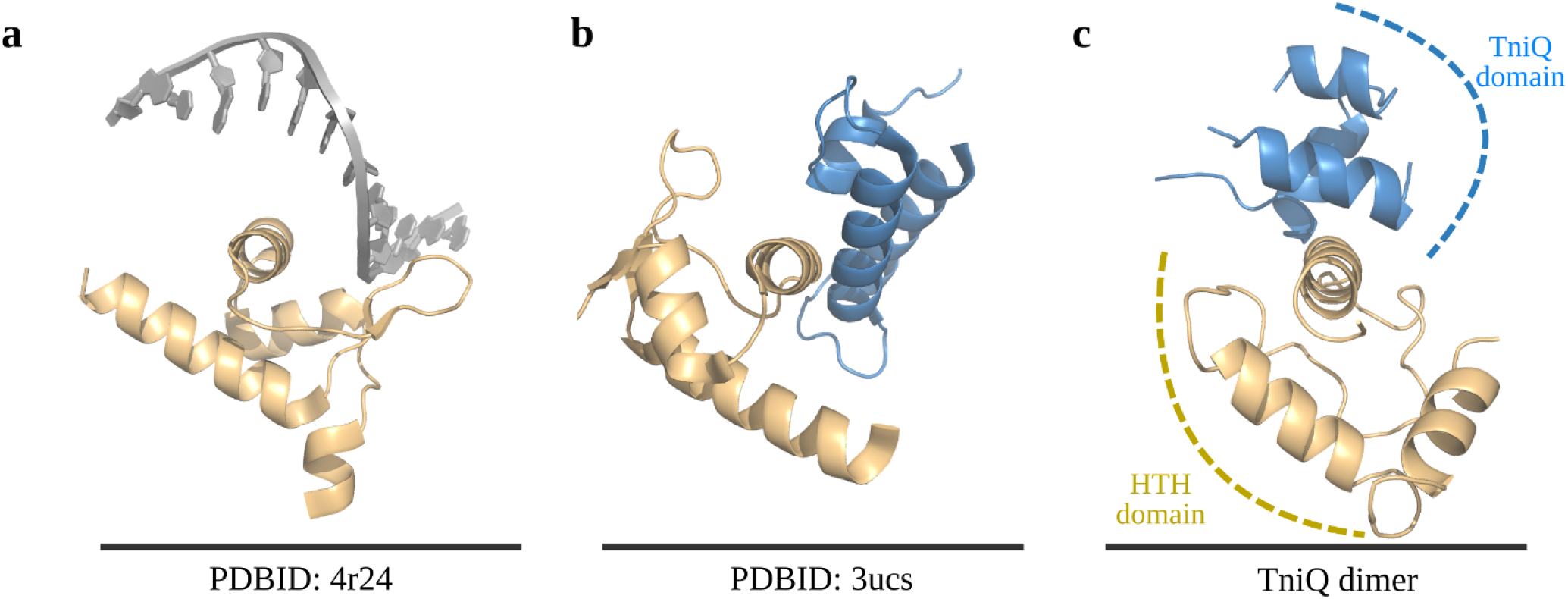
TniQ harbors a HTH domain involved in proteinprotein interactions within the TniQ dimer. A DALI search15 using the refined TniQ model as probe found significant similarity between the N-terminal domain of TniQ with PDB entries 4r24 (**a**) and 3ucs (**b**) (Z score 4.1/4.1, r.m.s.d. 3.8/5.1). Both proteins contain Helix-Turn-Helix (HTH) domains and HTH domains are often involved in nucleic acid recognition and mediate protein-protein interactions [15]. **c**, The TniQ dimer is stabilized in a head-to-tail configuration by reciprocal interactions mediated by the HTH domain and the TniQ-domains from both monomers.

**Extended Data Fig. 9.**
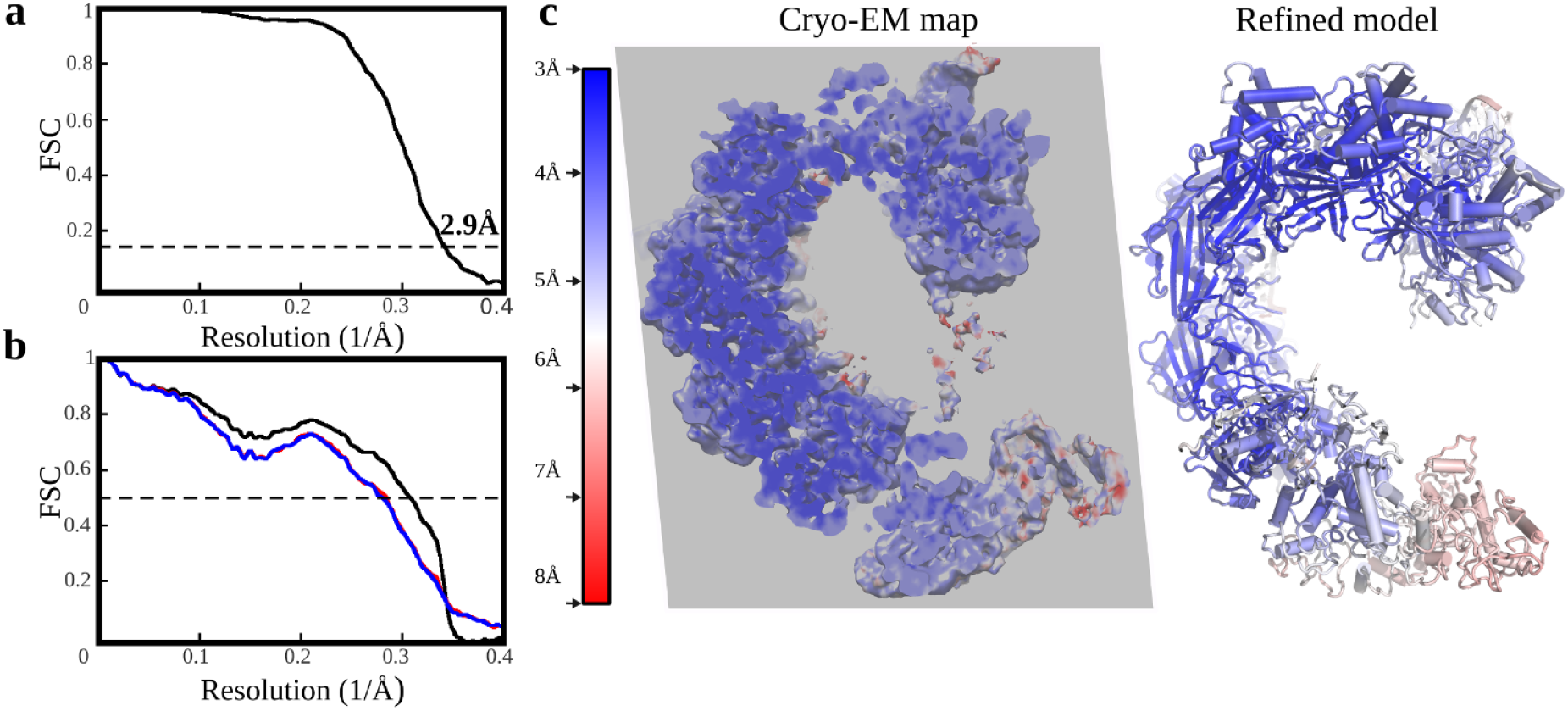
Fourier Shell Correlation (FSC) curves, local resolution, and unsharpened filter maps for the DNA-bound TniQ-Cascade complex complex. **a**, Gold-standard FSC curve using half maps; the global resolution estimation is 2.9 Å by the FSC 0.143 criterion. **b**, Cross-validation model-vs-map FSC. Blue curve, FSC between the shacked model refined against half map 1; red curve, FSC against half map 2, not included in the refinement; black curve, FSC between final model against the final map. The overlap observed between the blue and red curves guarantees a non-overfitted model. **c**, Left, unsharpened map colored according to local resolutions, as reported by RESMAP. dsDNA is visible at the top right projecting outside of the complex. Right, final model colored according to B-factors calculated by REFMAC.

**Extended Data Fig. 10.**
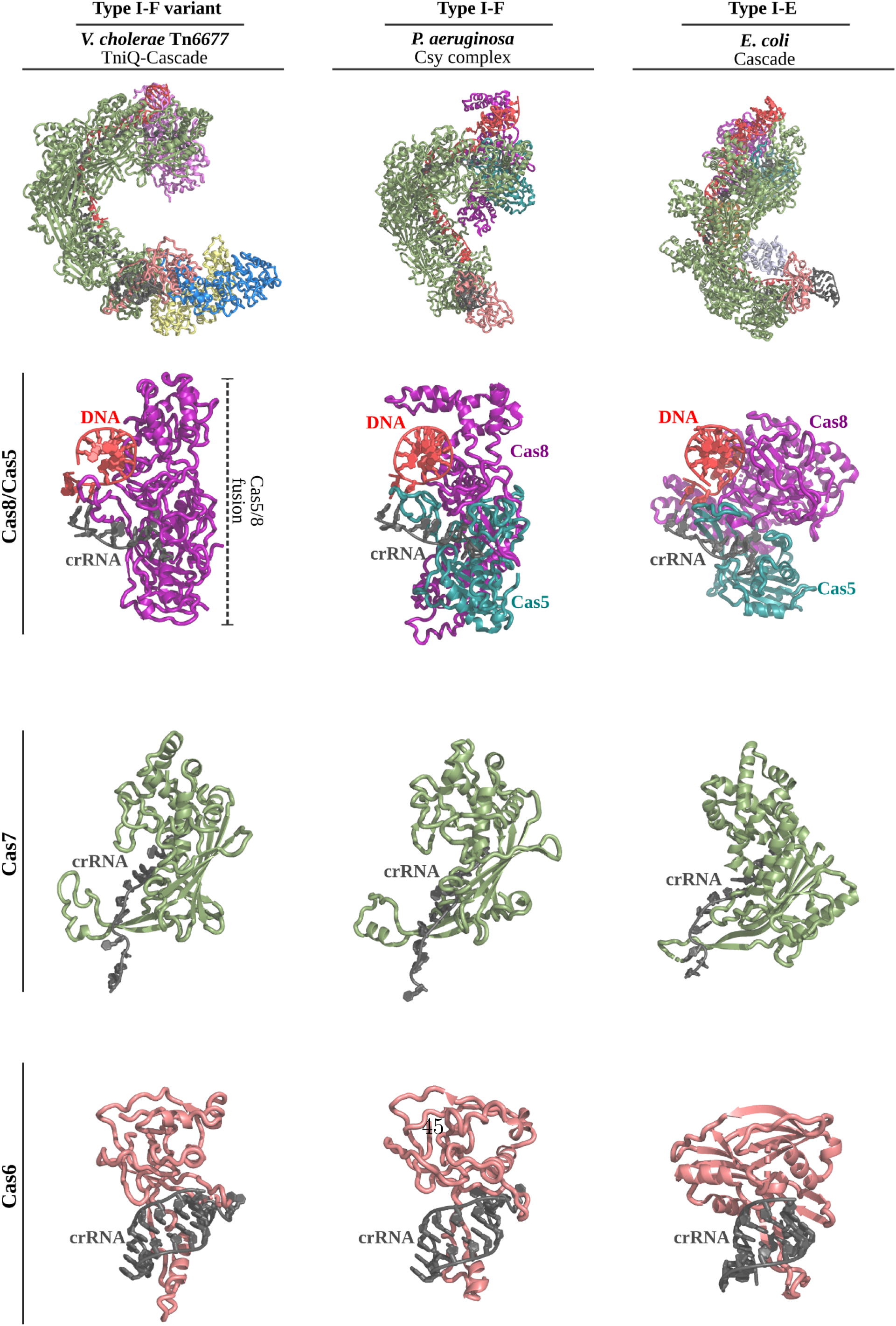
Superposition of DNA-bound TniQ-Cascade with structurally similar Cascade complexes. The DNA-bound structure of *V. cholerae* I-F variant TniQ-Cascade complex (left) was superposed with DNA-bound structures of *Pseudomonas aeruginosa* I-F Cascade11 (also known as Csy complex; middle, PDB ID: 6B44) and *Escherichia coli* I-E Cascade9 (right, PDB ID: 5H9F). Shown are superpositions of the entire complex (top), the Cas8 and Cas5 subunits with the 5’ crRNA handle and double-stranded PAM DNA (middle top), the Cas7 subunit with a fragment of crRNA (middle bottom), and the Cas6 subunit with the 3’ crRNA handle (bottom).

